# Activation of the innate immune system accelerates growth in cooperation with oncogenic Ras

**DOI:** 10.1101/2024.05.23.595555

**Authors:** Fabienne Brutscher, Federico Germani, George Hausmann, Lena Jutz, Konrad Basler

## Abstract

Innate immunity in *Drosophila* acts as an organismal surveillance system that measures external stimuli or cellular fitness and triggers context-specific responses to fight infections and maintain tissue homeostasis. However, uncontrolled activation of innate immune pathways can be detrimental to the system. In mammals, innate immune signaling is often overactivated in malignant cells and contributes to tumor progression. *Drosophila* tumor models have been instrumental in the discovery of interactions between pathways that promote tumorigenesis, but little is known about whether and how the Toll innate immune pathway interacts with oncogenes.

Here we use a *Drosophila* epithelial *in vivo* model to investigate the interplay between Toll signaling and oncogenic Ras. In absence of oncogenic Ras, Toll signaling suppresses differentiation and induces apoptosis. In contrast, in the context of Ras^V12^, cells are protected from cell death and Dorsal promotes cell survival and proliferation to drive hyperplasia. Taken together, we show that the tissue-protective functions of innate immune activity can be hijacked by pre-malignant cells to induce oncogenic transformation.

## Introduction

One of the main *Drosophila* innate immune signaling pathways is the Toll pathway (1). Toll signaling was initially discovered in the Nobel prize winning screen for genes involved in early *Drosophila* embryonic patterning (2). In subsequent research, Toll signaling was found to also be required for the innate immune response to infection (3). The signaling cascade is conserved from invertebrates to mammals and leads to the nuclear translocation of the NF-κB (nuclear factor ’kappa-light-chain-enhancer’ of activated B-cells)/ Rel-like (4) transcription factors, Dorsal and Dif. In the nucleus they regulate diverse genes including the production of antimicrobial peptides to attack invading pathogens (5,6).

The Toll pathway also acts as an organismal surveillance system that measures cellular fitness during development (7) and during cell competition (8,9). Less fit cells with elevated Toll activity are eliminated via apoptosis; this prevents abnormal cells from contributing to the structure of the tissue and thus helps maintaining tissue homeostasis (7–9).

In mammals, innate immunity provides the first line of defense against infection, but aberrant activation can be detrimental to the system (10). The NF-κB transcription factors of theToll-like receptor pathway (TLR; homologous to the *Drosophila* Toll pathway) are frequently overactivated in different malignant cells types and contribute to tumor progression by affecting processes such as cell survival and proliferation, invasion, metastasis and angiogenesis (11). Their role in cancer, whether it is pro- or anti-tumorigenic, appears to vary depending on the context (12,13). Ras is frequently mutated in cancer (14), but malignant transformation requires multiple steps of oncogene activation and inactivation of tumor suppressor genes (15). *Drosophila* tumor models have been instrumental in the discovery of factors involved in such cooperative oncogenesis as reviewed in (16,17). If and how the Toll pathway functions in the initial stages of Ras-induced oncogenesis needs further investigation.

Here we use a *Drosophila* epithelial model in the eye-antennal disc (EAD) to drive the expression of Ras^V12^, an oncogenic version of Ras and characterize the function of Toll pathway activation in this context. Interestingly, we found that in cells expressing Ras^V12^, Toll signaling potently drives tissue hyperplasia. To elucidate the mechanism underlying this effect, we investigated the transcriptional changes in Ras^V12^-transformed epithelia in response to Toll activation by constitutive expression of NF-κB/ Dorsal. During development, in the absence of oncogenic Ras, overexpression of Dorsal suppresses differentiation and induces apoptosis. In contrast, in the context of oncogenic Ras, cells are protected from cell death and Dorsal accelerates tumor growth. Overall, our results suggest that the tissue-protective functions of innate immune activity can be hijacked by premalignant cells to induce oncogenic transformation.

## Results

### Toll signaling cooperates with oncogenic Ras to drive overgrowth

To explore the role of TLR/ NF-κB signaling in the context of oncogene activation, we characterized the effects of Toll pathway activation in a widely used epithelial tumor model in the *Drosophila* larva (18–20): Using *eyeless* promoter-driven Flippase expression (*eyFlp*), we introduced genetic alterations, such as overexpression of an oncogene, into GFP-labelled cells in developing eye-antennal discs (EAD) (21). The size of EAD tumors was quantified at a late larval stage, at 96 h after egg deposition (AED) by three-dimensional reconstruction of GFP fluorescent signals from confocal image stacks. Consistent with a previous study (19), overexpression of a constitutive active form of Ras (Ras^V12^) alone impaired EAD development, but did not induce hyperplastic growth (Fig 1A and 1B). Next, we examined the effect of altering Toll signaling. In the absence of Ras^V12^, overexpression of the NF-κB/ Rel-like transcription factor Dorsal impaired EAD development and reduced EAD size (Fig 1A and 1B). Interestingly, in combination with Ras^V12^, activation of the Toll pathway through overexpression of Dorsal induced overgrowth (Fig 1A and 1B). The same result was seen when overexpressing a ligand-independent, constitutively active Toll receptor (Toll^10b^) (22) to activate Toll signaling (Fig 1A and 1B). Inhibition of Toll signaling by overexpressing Cactus, the inhibitor of Dorsal (homologous to mammalian Inhibitor of κB (IκB)) appears to have a slight tumor suppressive function in combination with Ras^V12^ (p=0.06) (Fig 1A and 1B). However, it should be noted that overexpression of Cactus also reduced the size of control EADs (Fig. 1A and 1B); This may have indirectly influenced tumor growth also in the context of Ras^V12^.

**Fig 1.**
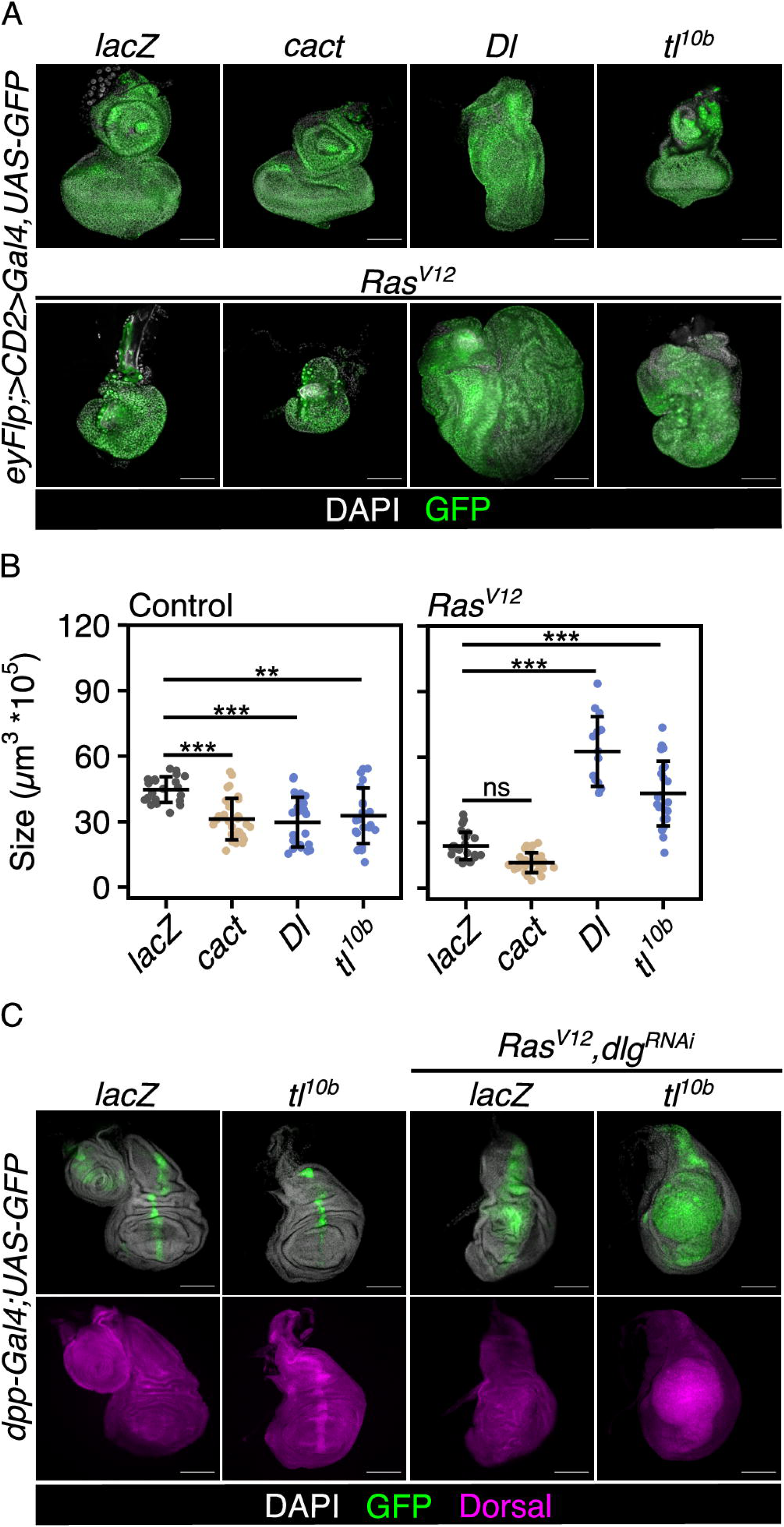
Toll signaling cooperates with oncogenic Ras to induce overgrowth. (A) Confocal images of EADs and EAD tumors at 96 h AED of the indicated genotypes representative for tissue sizes quantified in (B). (B) Quantification of tissue volume of the indicated genotypes. Mean volume and standard deviation (error bar) are shown (***p < 0.001, **p < 0.01, ns: p = 0.06, One-way ANOVA with post-hoc Tukey HSD). (C) Confocal images of wing discs of indicated genotypes at 120 h AED labelled for Dorsal (magenta) as a read-out for Toll pathway activity. In all confocal images DAPI (grey) is used to visualize nuclei and GFP (green) labels cells co-expressing indicated UAS-transgenes. Scale bars represent 100 µm. ns: not significant, EAD: eye-antennal disc, AED: after egg deposition, cact: Cactus, Dl: Dorsal, tl^10b^: Toll10b, eyFlp: eyeless-driven Flippase, dpp: Decapentaplegic.

Toll signaling also enhanced the overgrowth of another neoplastic tumor generated by impairing apical-basal cell polarity by knockdown of Disc large (dlg) and simultaneously overexpressing Ras^V12^ (Ras^V12^, dlg^RNAi^) (23) (Fig S1A and S1B). The effect of Dorsal overexpression on the size of Ras^V12^, dlg^RNAi^, however, was only apparent later during tumor development, at 120 h AED compared to the 96 h AED effect in the context of Ras^V12^ alone (Fig S1B and S1C). The size of EADs expressing only dlg^RNAi^ was unaffected by Dorsal overexpression (Fig S1D and S1E).

Inhibition of Toll signaling by overexpressing Cactus did not affect the size of Ras^V12^, dlg^RNAi^-induced tumors (Fig S1A and S1B). However, overexpression of Cactus was able to rescue the morphological defects caused by Toll activation in control EADs (Fig S1F). Additionally, Dorsal was not enriched in Ras^V12^, dlg^RNAi^-tumors induced in the Dpp (Decapentaplegic) stripe of the wing disc (Fig 1C). Hence, we conclude that Toll signaling was not activated by nor required for Ras^V12^, dlg^RNAi^ driven tumor growth, but able to promote overgrowth if ectopically activated. How activation of the innate immune system interacts with oncogenes to promote overgrowth is unclear.

### Dorsal inhibits retinal differentiation

The overgrowth of Ras^V12^-transformed EADs after Toll pathway activation could have various explanations, including effects on differentiation, proliferation and apoptosis. To explore which of these processes could have contributed to the overgrowth in Ras^V12^, Dorsal tumors, we first investigated the transcriptional changes in Ras^V12^-expressing EADs in response to Toll activation. By choosing a non-clonal induction system (23), we were able to identify transcriptional changes in a homogeneously transformed cell population, circumventing potential confounding non-autonomous effects from wild-type cells. We sequenced the transcriptomes of control and Ras^V12^-transformed EADs with and without Toll pathway activation induced by Dorsal overexpression. A high number of genes was significantly differentially expressed in each of the comparisons (Fig 2A). Confirming we had activated the Toll pathway, when comparing Ras^V12^, Dorsal to Ras^V12^, lacZ tumors, we found many innate immune pathway associated genes, such as *cactus* (*cact*), *Peptidoglycan recognition protein SA* (*PGRP-SA*) or *wnt inhibitor of Dorsal* (*wntD*), amongst the most upregulated genes (Fig 2B and S2A).

**Fig 2.**
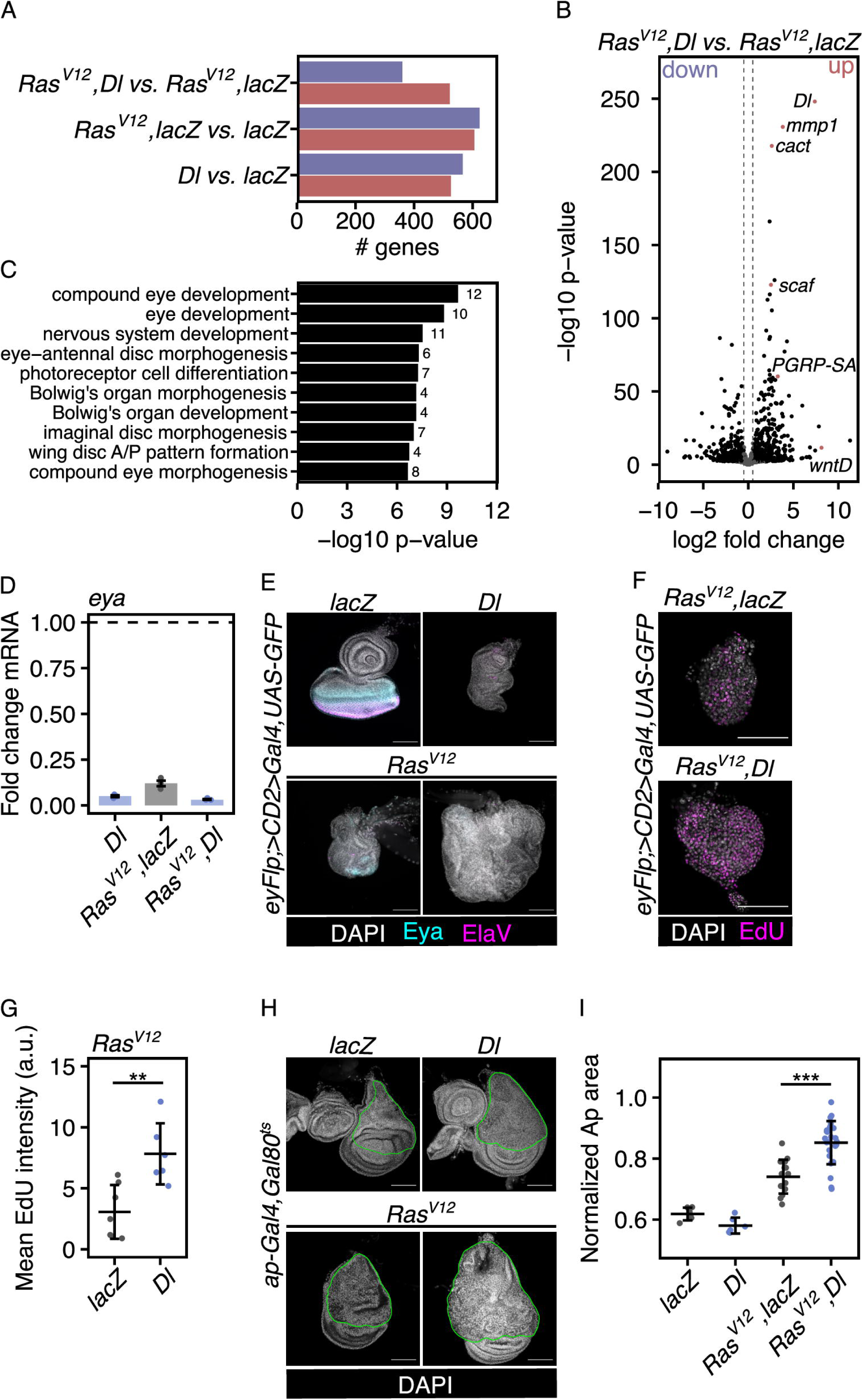
Dorsal inhibits retinal differentiation. (A) Total number and directionality (blue: downregulated, red: upregulated) of significantly differentially expressed genes between indicated genotypes illustrating similarity relative to each other. (B) Volcano plot of genes significantly differentially expressed between Ras^V12^, Dorsal and Ras^V12^, lacZ tumors. Significantly differentially expressed genes (p < 0.05, absolute log2 fold > 0.5) are shown in black and non-significant genes in grey. A positive fold change indicates higher expression in Ras^V12^, Dorsal compared to Ras^V12^, lacZ tumors. Vertical lines indicate the absolute log2 fold thresholdsof 0.5 and -0.5. Exemplary upregulated genes of interest are annotated and highlighted in red. (C) Bar chart of the ten most enriched gene sets for the expression signature of genes downregulated after Dorsal overexpression identified through hypergeometric Gene Set Enrichment Analysis (GSEA) on the gene set databases Gene Ontology, Biological Process (GO:BP). Data labels represent the number of significantly differentially expressed genes within each gene set. Significantly enriched gene sets (p < 0.05, absolute log2 fold > 0.0) are depicted in black. (D) qRT-PCR analysis of *eya* mRNA levels in EADs expressing indicated UAS-transgenes under the control of an eye specific Gal4 (*eyFlp; act>CD2>Gal4*) as exemplary confirmation of the downregulation of a retinal determination gene after overexpression of Dorsal. Data is shown as fold changes relative to the control (lacZ). n=3 biologically independent samples were analysed. Median fold changes and interquartile ranges (error bars) are shown. The dashed horizontal line illustrates the reference fold change of 1. (E) Confocal images of EADs and EAD tumors of the indicated genotypes at 96 h AED labelled for retinal determination and photoreceptor differentiation markers Eya (cyan) and ElaV (magenta). (F) Confocal images of EAD tumors of indicated genotypes at 72 h AED stained with Click-iT™ EdU (5-ethynyl-2’-deoxyuridine) Alexa Fluor™ 647 (magenta) to identify DNA synthesis-based proliferation, representative for EdU intensity quantified in (G); scale bars represent 20 µm. (G) Quantification of EdU mean fluorescence intensity per EAD tumor at 72 h AED. Mean volume and standard deviation (error bar) are shown (**p < 0.01, One-way ANOVA with post-hoc Tukey HSD). (H) Confocal images of wing discs of the indicated genotypes representative for relative sizes quantified in (I). The green outline is based on the pattern of Gal4 immunoreactivity and highlights the dorsal, Apterous-marked compartment. (I) Quantification of the Apterous-marked area relative to overall disc area for indicated genotypes. Mean area and standard deviation (error bar) are shown (***p < 0.001, One-way ANOVA with post-hoc Tukey HSD). In all confocal images DAPI (grey) is used to visualize nuclei and GFP (green) labels cells co-expressing indicated UAS-transgenes. Scale bars represent 100 µm, unless otherwise indicated. a.u.: arbitrary unit. EAD: eye-antennal disc, AED: after egg deposition, Dl: Dorsal, wntD: wnt inhibitor of Dorsal, PGRP-SA: Peptidoglycan recognition protein SA, scaf: scarface, mmp1: matrix metalloproteinase 1, eyFlp: eyeless-driven Flippase, ap: Apterous, Eya: eyes absent, ElaV: embryonic lethal abnormal vision.

Interestingly, overexpression of Dorsal also affected differentiation. In both the control and Ras^V12^ context, Dorsal overexpression resulted in the significant downregulation of genes involved in processes connected to EAD development (Fig 2C). Consistent with an impaired EAD development we found that markers for retinal determination and photoreceptor differentiation, such as Eyes absent (Eya) and Embryonic lethal abnormal vision (ElaV), were downregulated after the overexpression of Dorsal (Fig 2D and E).

These results could suggest that Dorsal overexpression traps cells in a progenitor-like state, maintaining a proliferative potential throughout the entire EAD. To explore this, we looked at proliferation at various stages. Overexpression of Dorsal in Ras^V12^-transformed EADs did increase proliferation. The increase was most pronounced during early tumor development, at 72 h AED (Fig 2F and 2G, Fig S2B and S2C), when both tumors (Ras^V12^, lacZ and Ras^V12^, Dorsal) were still of the same size (Fig S2D). Intriguingly, the overexpression of Toll^10b^ (Tl^10b^), although it was able to induce overgrowth in the context of Ras^V12^ (Fig 1A and B), did not noticeably suppress retinal determination and photoreceptor differentiation (Fig S2E). Thus, while inhibition of retinal differentiation likely contributes to overgrowth induced by Toll activation, it is not necessary.

To further test if an inhibition of differentiation is required for enhanced tumor growth by Toll activation, we examined the consequences of co-overexpressing Ras^V12^ and Dorsal in the dorsal compartment of the wing disc. During the experimental time window, the cells of the dorsal compartment are not yet differentiated (24). We found that growth of the dorsal compartment, marked by the expression of Apterous, was enhanced when Dorsal was overexpressed together with Ras^V12^ (Fig2H and 2I). This demonstrated that the growth-promoting effect of Toll signaling on Ras^V12^-driven tumors is not limited to the EAD and can occur independently of the differentiation status.

### Dorsal promotes overgrowth independent of JNK signaling

Among the genes differentially expressed following Toll activation were various targets of the JNK pathway. Previous reports have demonstrated that Toll signaling can activate the JNK pathway (25–27). To investigate a possible contribution of JNK to the Toll signaling-induced overgrowth in our system, we analyzed the transcriptional changes after Toll activation that are associated with the JNK pathway. Many genes associated with increased JNK signaling were strongly upregulated after Dorsal overexpression (Fig 3A). For example, the expression of JNK-regulated genes, such as *scarface* (*scaf*) (28), *matrix metalloproteinase 1* (*mmp1*) (29–31) and *Ets at 21c* (*Ets21c*) (32,33) was elevated in Ras^V12^, Dorsal tumors (Fig 3A, 3B and 3C, Fig S3A).

**Fig 3.**
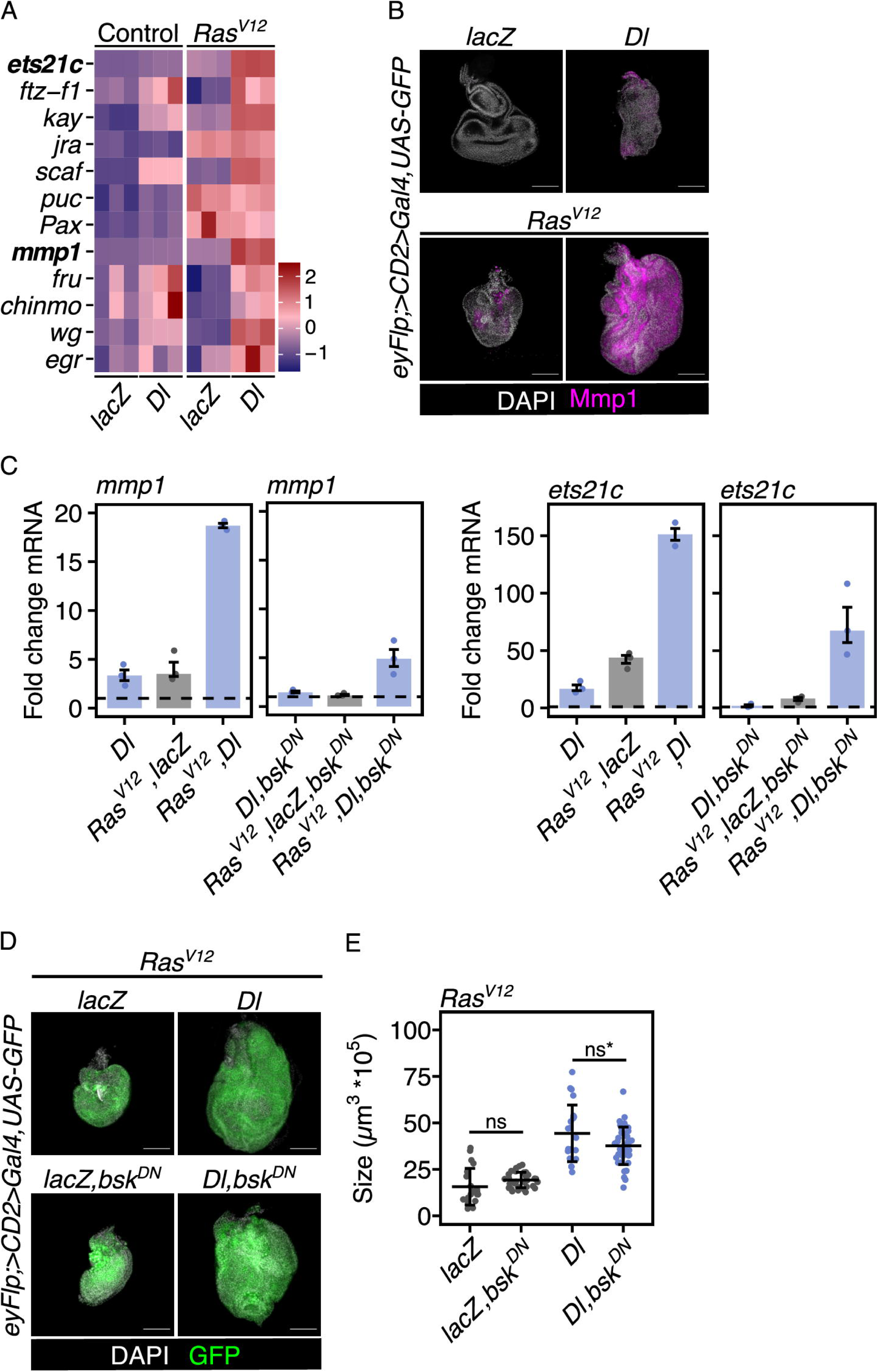
Dorsal promotes overgrowth independent of JNK signaling. (A) Heatmap showing the expression pattern of genes commonly associated with JNK signaling across all four samples. Colour intensity represents the expression level (lowest to highest from blue to red) scaled into z-scores. Two genes of interest, *ets21c* and *mmp1*, are depicted in bold. (B) Confocal images of EADs and EAD tumors of indicated genotypes at 96 h AED, labelled for Mmp1 confirming the upregulation of one of the JNK-regulated genes shown in (A). (C) qRT-PCR analysis of *mmp1* and *ets21c* mRNA levels in EADs expressing indicated UAS-transgenes under the control of an eye specific Gal4 (*eyFlp; act>CD2>Gal4*) as exemplary confirmation of the JNK-dependent upregulation of target genes after overexpression of Dorsal. Data is shown as fold changes relative to the control (lacZ or lacZ,bsk^DN^). n=3 biologically independent samples were analysed. Median fold changes and interquartile ranges (error bars) are shown. The dashed horizontal line illustrates the reference fold change of 1. (D) Confocal images of EADs tumors of indicated genotypes at 96 h AED representative for tissue sizes quantified in (E). (E) Quantification of tissue volume of indicated genotypes at 96 h AED before and after JNK pathway inhibition. Mean volume and standard deviation (error bar) are shown (ns: p=0.6, ns*:p= 0.07), One-way ANOVA with post-hoc Tukey HSD). In all confocal images DAPI (grey) is used to visualize nuclei and GFP (green) labels cells co-expressing indicated UAS-transgenes. Scale bars represent 100 µm. ns: not significant, EAD: eye-antennal disc, AED: after egg deposition, Dl: Dorsal, mmp1: matrix metalloproteinase 1, ets21c: Ets at 21c, bsk^DN^: dominant-negative Basket, JNK: Jun-N-terminal Kinase pathway, eyFlp: eyeless-driven Flippase.

Previous studies had suggested that JNK is required for malignancy (invasiveness), but not overgrowth of *Ras^V12^,scribble* mutant tumors (18,30). To inhibit JNK signaling in our system we either co-expressed a dominant-negative form of the *Drosophila* JNK Basket (Bsk^DN^) or the negative JNK pathway regulator Puckered (Puc). Confirming we had effectively inhibited the pathway, the transcription of JNK target genes was markedly decreased by these manipulations (Fig 3C and S3A). Consistent with the notion that JNK drives malignancy but not overgrowth, inhibiting JNK in Ras^V12^, Dorsal tumors did not markedly reduce tissue overgrowth (Fig 3D and 3E, Fig S3C). Also, the increase in size of Ras^V12^, dlg^RNAi^ tumors caused by the overexpression of Dorsal was independent of JNK signaling (Fig S3D). Taken together, these results suggest that factors other than enhanced JNK signaling can mediate the growth stimulatory effect that Toll activation exerts on Ras^V12^ signalling.

### Dorsal-induced caspase-dependent cell death is suppressed by Ras^V12^

We found that Dorsal affected differentiation and proliferation in Ras^V12^-transformed EADs. A third feature that could have contributed to the increased growth of these tumors is changes in cell death rate. Therefore, we asked if apoptosis levels changed in the different conditions. In EADs, in the absence of Ras^V12^, the levels of activated effector caspase, cleaved *Drosophila* caspase-1 (Dcp1), were strongly elevated in response to Toll activation (Fig 4A and 4B). Consistent with this, the expression of the pro-apoptotic gene, *reaper,* was upregulated (Fig 4C). These effects were independent of JNK signaling as expressing Bsk^DN^ did not abolish either Dcp1 or *rpr* expression (Fig S4A, S4B and S4C). In the context of Ras^V12^, Dorsal-induced cell death was inhibited: The mRNA levels of Dcp1 and Reaper were significantly lower compared to those when Dorsal was overexpressed in control EADs (Fig 4A, 4B and 4C).

**Fig 4.**
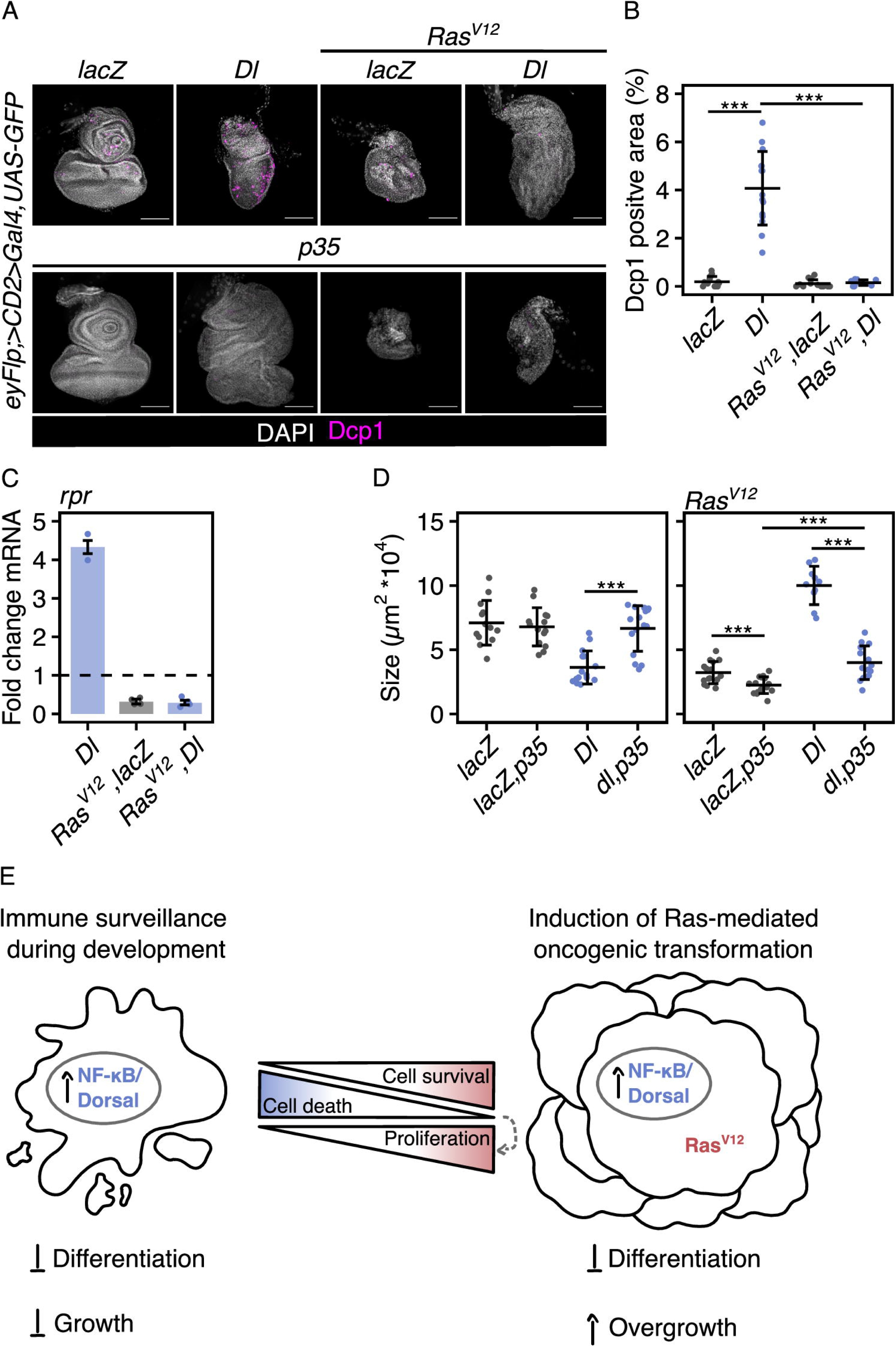
Dorsal-induced caspase-dependent cell death is suppressed by Ras^V12^. (A) Confocal images of EADs and tumors of indicated genotypes at 96 h AED labelled for Dcp1 illustrating the enrichment of cell death after Dorsal overexpression in control EADs as quantified in (B) and demonstrating the loss of cell death after inhibition of apoptosis through overexpression of p35. (B) Quantification of the Dcp1 positive area relative to whole disc size at 96 h AED. Mean area and standard deviation (error bar) are shown (***p < 0.001, One-way ANOVA with post-hoc Tukey HSD). (C) qRT-PCR analysis of *rpr* mRNA levels in EADs expressing indicated UAS-transgenes under the control of an eye specific Gal4 (*eyFlp; act>CD2>Gal4*) illustrating the increase in pro-apoptotic gene expression after overexpression of Dorsal in control EADs. Data is shown as fold changes relative to the control (lacZ). n=3 or n=4 biologically independent samples were analysed. Median fold changes and interquartile ranges (error bars) are shown. The dashed horizontal line illustrates the reference fold change of 1. (D) Quantification of tissue size of indicated genotypes at 96 h AED to illustrate changes to the effects of Dorsal on tissue growth after inhibition of apoptosis through co-expression of p35. Mean area and standard deviation (error bar) are shown (***p < 0.001, One-way ANOVA with post-hoc Tukey HSD). (E) Graphical model illustrating the transition of the function of NF-κB/ Dorsal from growth-suppressive during development to growth-promoting during Ras^V12^-mediated oncogenesis. The transition is mediated by the combined effects of Dorsal-induced changes in cell death, cell survival, proliferation and differentiation. In all confocal images DAPI (grey) is used to visualize nuclei and GFP (green) labels cells co-expressing indicated UAS-transgenes. Scale bars represent 100 µm. EAD: eye-antennal disc, AED: after egg deposition, Dl: Dorsal, Dcp1: cleaved *Drosophila* caspase 1, rpr: reaper, NF-κB: nuclear factor ’kappa-light-chain-enhancer’ of activated B-cells.

The inhibition of cell death mediated by Ras^V12^ may enable Dorsal to promote tissue overgrowth. We therefore next asked how specifically inhibiting apoptosis in the context of Dorsal overexpression affected tissue growth. To this end we co-expressed the baculoviral anti-apoptosis protein p35, an inhibitor of effector caspases, such as Dcp1 (34,35). Co-expression of p35 prevented the caspase activation seen in response to overexpressing Dorsal: no active Dcp1 was detected (Fig 4A). As expected, EAD size was increased to the level of control EAD size (Fig 4D). Since Dorsal still impaired retinal differentiation, the EAD was malformed (Fig S4D).

Interestingly, in the context of Ras^V12^, p35-mediated inhibition of apoptosis further affected tissue growth (Fig 4A and 4D). Indeed, p35 co-expression with Ras^V12^ seemed to impair growth and to reduce tumor size. The growth enhancement induced by Dorsal overexpression was however not abolished by co-expression of p35: Ras^V12^, Dorsal, p35 EADs were still larger than Ras^V12^, lacZ, p35 EADs (Fig 4D and S4E). Based on the reduced *rpr* expression and on the absence of detectable levels of Dcp1, we contend that Ras^V12^ can prevent programmed cell death, thus driving Dorsal-induced overgrowth.

## Discussion

Using *Drosophila* imaginal epithelia as a model we have explored how the Toll pathway cooperates with oncogenic Ras to drive tumor growth. Although Toll activation restricts growth during EAD development, we found that when activated in the context of oncogenic Ras, Toll signaling induces overgrowth. We then evaluated three potential explanations for the enhanced growth: differentiation, proliferation and apoptosis.

Based on a transcriptome analysis we found that the activation of the Toll pathway impaired retinal differentiation, trapping cells in a progenitor-like state (Fig 2). Increased cell death rates prevent EADs from overgrowing after Toll activation (Fig 3). However, these cells maintain a proliferative potential throughout the entire EAD. Hence, overexpressing Dorsal in the context of Ras^V12^, which renders cells resistant to cell death, results in increased proliferation and accelerated tissue growth. However, two lines of evidence indicated that impaired differentiation in combination with Ras^V12^ is not a prerequisite to induce tissue overgrowth: (1) Activation of the Toll pathway via Toll^10b^ induced overgrowth even though it did not impair differentiation. (2) In the WD, in which cells do not differentiate during the experimental time window, activating Toll signaling also induced overgrowth in cooperation with Ras^V12^.

We also found that increased proliferation was most pronounced early after tumor induction, demonstrating the importance of careful assessment of tumor development over multiple time points.

Consistent with previous reports in *Drosophila* (7,8,26,27,36), we found that Dorsal activation is pro-apoptotic and induces cell death during development (Fig4 and S4). In the context of oncogenic Ras, however, apoptosis is suppressed (37,38), and the Dorsal-mediated outputs were tumor-promoting. The shift from anti-to pro-tumorigenic function is often associated with the ambivalent nature of JNK signaling (39). This has, for example, been demonstrated in the context of polarity-deficient (*scribble* mutant) cell clones co-expressing Ras^V12^. In this context, rather than promoting apoptosis, JNK has a pro-tumorigenic function and drives tumor cell invasion (30,31,40–42). JNK has been reported to be a target of the Toll pathway in different contexts (10,11,14). It was thus not unexpected that we found JNK signaling upregulated upon Dorsal overexpression. However, blocking JNK signaling did not significantly affect Dorsal-induced overgrowth in the context of Ras^V12^ (Fig 2-4 and S2-4). As the expression of the common JNK target genes, *mmp1* and *ets21c*, was not completely suppressed despite JNK inhibition (Fig 3C and S3A), we cannot exclude that they continue contributing to the Dorsal-induced effects. Especially, the transcription factor Ets21c was previously shown to drive tumor growth downstream of JNK in cooperation with Ras^V12^ and is predicted to be regulated by inputs in addition to that of JNK (17,18). Ets21c may thus still have contributed to the observed overgrowth of Ras^V12^, Dorsal discs.

The inhibition of cell death by oncogenic Ras may have enabled Dorsal to switch from limiting tissue growth by inducing apoptosis to promoting overgrowth by enhancing proliferation. To induce overgrowth, Dorsal may directly regulate the expression of genes that promote apoptosis and proliferation. While in control EADs the induction of apoptosis may outweigh a potential increase in proliferation, the positive effects of Dorsal on proliferation and tissue growth prevail when cell death is inhibited, e.g. via Ras^V12^. Caspases are known to not only drive cell death, but they can also have non-apoptotic functions (43), such as caspase-induced proliferation leading to tissue hyperplasia (44,45). We therefore suggest that in the context of Ras^V12^, in which apoptosis is suppressed, Toll signaling may also promote overgrowth through caspase-induced proliferation. Taken together, our data indicates that Toll/ Dorsal accelerates tumor growth in cooperation with oncogenic Ras likely through the combined effects on differentiation and apoptosis.

Even though mammalian TLR/NF-κB signaling is pre-dominantly considered a tumor promoting pathway (46), in certain contexts, NF-κB can act as a tumor-suppressor (47–51). Currently, to the best of our knowledge, *in vivo* studies investigating the relationship between tumor-intrinsic oncogenic Ras and NF-κB are rare (52,53). With the present study, we used the *Drosophila* larva as a simple *in vivo* system to investigate the interplay between NF-κB/ Dorsal and oncogenic Ras. In the EAD we found that activation of Toll signaling suppressed retinal differentiation and induced caspase-activation. In the context of oncogenic Ras, these consequences likely contribute to tumorous overgrowth.

## Materials and methods

### *Drosophila* stocks and husbandry

Flies were reared and maintained at 25°C (12:12 h light/dark cycle) on standard corn-meal food containing, per liter, 100 g fresh yeast, 55 g corn meal powder, 10 g organic wheat flour, 8 g agar, 75 g white sugar and 15 ml nipagin. The fly strains used in this study and listed in Table 1 were mainly obtained from the Bloomington *Drosophila* Stock Center (BDSC), the Vienna *Drosophila* Resource Center (VDRC) or, for UAS-ORF constructs inserted at position 86FB (54), from the FlyORF Injection Service. Crosses were raised at 25°C, except those involving *tub-Gal80^ts^*, which were raised at 18°C and moved to 29°C, according to the experimental set up. To induce the expression of transgenes in the EAD, virgin flies of *eyFlp; ; act>CD2>Gal4,UAS-GFP* or *eyFlp; UAS-Ras^V12^,UAS-dlg^RNAi^/CyO,tub-Gal80; act* > *CD2* > *Gal4, UAS-GFP* were crossed to respective *UAS*-constructs. To induce transgene expression in the WD, virgin flies of *dpp-Gal4* or *ap-Gal4,tub-Gal80ts* were crossed to respective *UAS*-constructs.

**Table 1.** Drosophila stocks used in this study.

| Genotype | Source | ID |
| --- | --- | --- |
| *ap-Gal4* | BDSC | 3041 |
| *dpp-Gal4* | n/a | n/a |
| *eyFlp1*; ; *act>CD2>Gal4,UAS-GFP<sup>S65T</sup>* | (23) | n/a |
| *GMR-Gal4* | (55) | n/a |
| *tub-Gal80* | BDSC | 9491 |
| *tub-Gal80<sup>ts</sup>* | BDSC | 7019 |
| *UAS-bsk<sup>DN</sup>* | Basler lab | n/a |
| *UAS-cact* | FlyORF | F001339 |
| *UAS-dl* | BDSC | 9319 |
| *UAS-dlg<sup>RNAi</sup>* | VDRC | 41134 |
| *UAS-egr* | (56) | n/a |
| *UAS-lacZ* | Basler lab | n/a |
| *UAS-p35* | BDSC | 5073 |
| *UAS-puc* | BDSC | 98328 |
| *UAS-Ras<sup>V12</sup>* | BDSC | 64196 |
| *UAS-*tl10b** | Basler lab | n/a |

### Immunohistochemistry

Micro-dissected imaginal disc tissues were fixed in 4 % formaldehyde (Table 4) in 1 x phosphate-buffered saline (PBS) for 20 min and washed three times for 5 min in PBS at room temperature (RT). Samples were permeabilized for 10 min in 0.1 % Triton X-100 in 1 x PBS (PBT) and blocked for 30 min in 2 % HINGS. Samples were incubated with primary antibody solutions listed in Table 2 at 4 °C overnight, washed three times for 5 min in PBS and then incubated with host-specific Alexa Fluor secondary antibodies (Alexa Fluor 555, 647) and DAPI (1:200) at RT. The samples were washed three times for 5 min before mounting in Vectashield (Table 4). To reduce tissue compression, for volume measurements, all EADs and EAD tumors were mounted on Poly-L-lysine coated glass bottom dishes (Table 4) without coverslips. Click-iT Plus EdU Alexa Fluor 647 Imaging Kit (Table 4) was used to quantify DNA synthesis (S-Phase) *in vivo*, following the manufacture’s protocol. EADs and EAD tumors were incubated in EdU labeling solution for 45 min at RT.

**Table 2.** Primary antibodies used in this study.

| Name | Source | Antibody registry ID | Dilution (in 1 x PBS) |
| --- | --- | --- | --- |
| mouse monoclonal anti-Dorsal | DSHB | AB_528204 | 1:20 |
| mouse monoclonal anti-Eya | DSHB | AB_528232 | 1:20 |
| mouse monoclonal anti-Mmp1 | DSHB | AB_579781 | 1:400 |
| rabbit polyclonal anti-cleaved Dcp1 | Cell Signaling Technologies | AB_2721060 | 1:200 |
| rabbit polyclonal anti-Gal4 (DBD): sc-577 | Santa Cruz Biotechnology | n/a | 1:1000 |
| rat monoclonal anti-Elav | DSHB | AB_528218 | 1:200 |

### Microscopy

Confocal images were acquired using an inverted laser scanning confocal microscope (Leica SP8 Inverse, HC PL APO CS2 20 x/0.75 NA objectives, 405 nm (50 mW), 488 nm (20 mW), 552 nm (20 mW) and 638 nm (30 mW) lasers). Single confocal planes were imaged and processed using the Fiji package of ImageJ (57). Brightfield images of adult eyes were acquired on Axio Zoom V16 (Zeiss) and were processed using Fiji.

### Quantification and statistical analysis of tumor growth

Tumor- and control EAD bearing larvae were dissected 72, 96 or 120 h AED as indicated for each experiment. Typically, we measured the size of tumors and control EADs using three-dimensional (3D) reconstruction of GFP-labeled tissue from confocal image stacks as previously described in (58). The confocal planes were imaged with the 10 x/0.3 NA objective (Leica Sp8 Inverse, HC PL APO) at 15 µm intervals until the entire sample was captured from top to bottom. 3D reconstruction and volume measurements were done using IMARIS image analysis software (Bitplane). To compare the sizes of control and tumor EADs with and without co-expression of p35, two-dimensional measurements were done using FIJI’s polygon tool. Tumor- and control WD bearing larvae were raised and maintained at 18 °C until moved to 29 °C three days AED (approximate equivalent developmental age at) until dissected three days later Tumor- and control area in the dorsal compartment of the WD was measured based on the region marked by Apterous using FIJI’s polygon tool. Data visualization and statistical analysis were carried out using R, version 4.2.3 (59). One-way ANOVA (analysis of variance) with post-hoc Tukey HSD (honestly significant difference) was used to compare the mean value between multiple groups.

### RNA isolation, cDNA synthesis and quantitative real-time PCR

Total RNA was extracted from approximately 50 control or 30 tumor EADs per sample from larvae of the respective genotypes at 96 h AED. Per genotype three biologically independent replicates were sampled. The samples were then snap-frozen in dry ice. RNA was isolated using the RNeasy Micro Kit (Table 4) following the manufacture’s protocol. cDNA was synthesized using the PrimeScript™ RT Master Mix (Table 4). Quantitative real-time PCR reactions were performed in technical triplicates using the PowerUp SYBR Master Mix (Table 4) and analysed using the QuantStudio3 system (Applied Biosystems). The sequences of the primers used in this study are listed in Table 3.

**Table 3.**
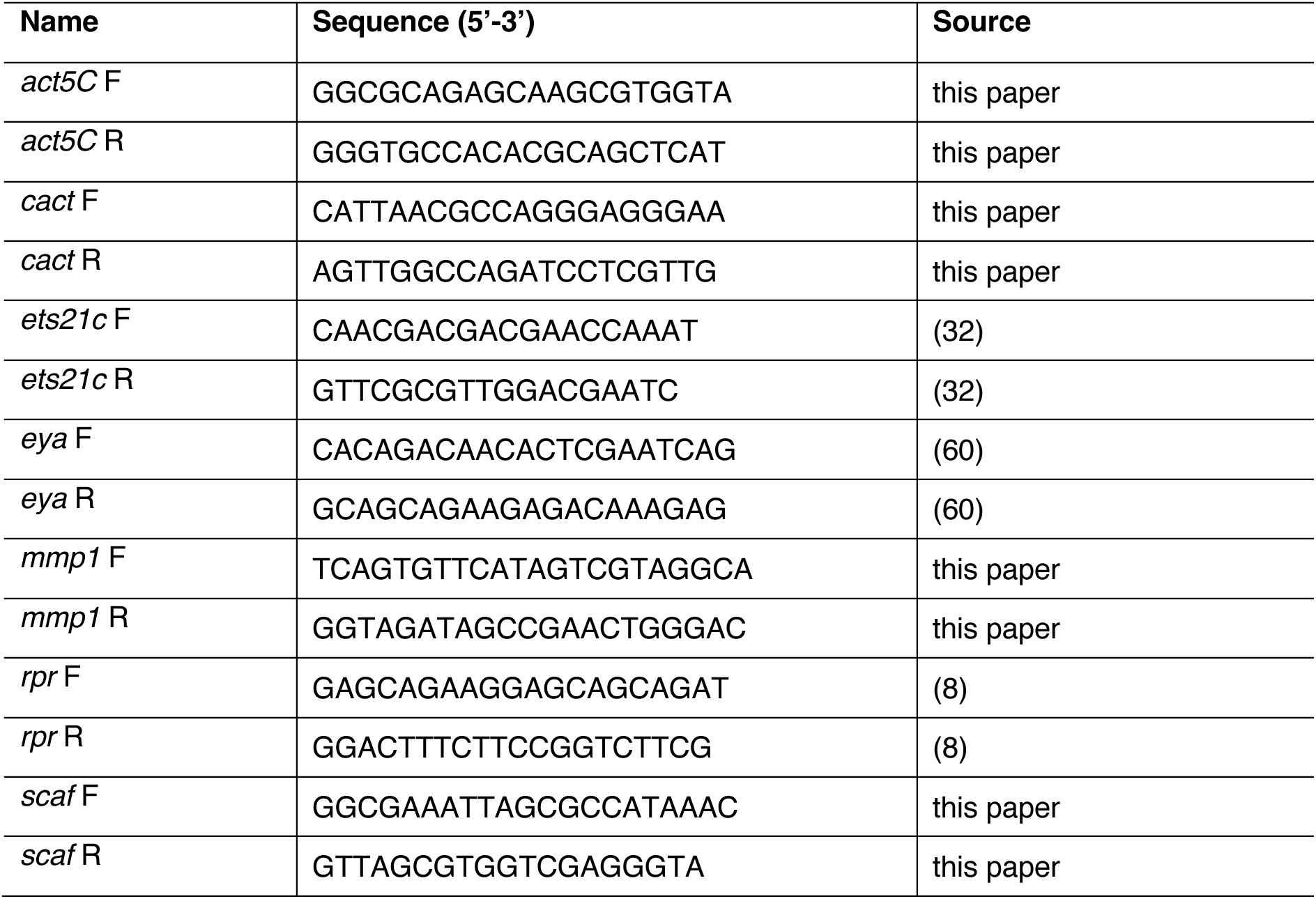
qRT-PCR primers used in this study.

**Table 4.**
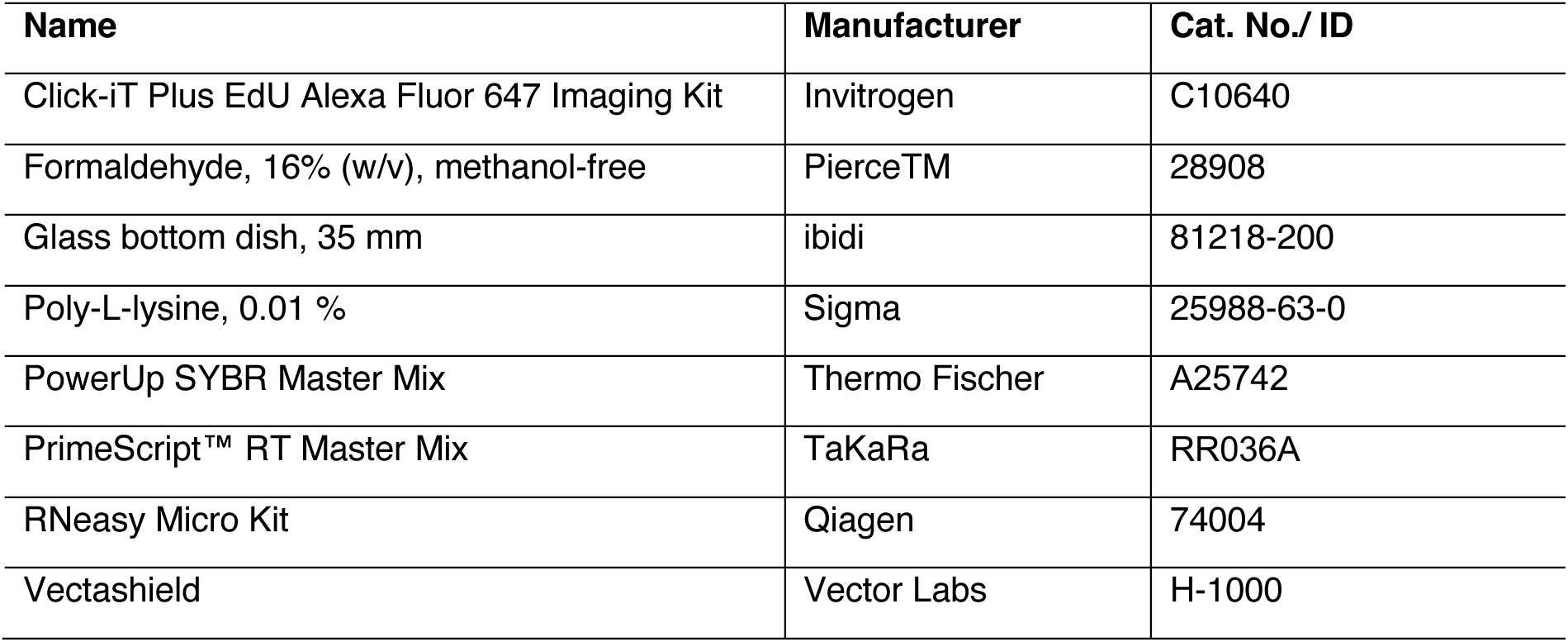
Extended list of materials used in this study.

| Name | Manufacturer | Cat. No./ ID |
| --- | --- | --- |
| Click-iT Plus EdU Alexa Fluor 647 Imaging Kit | Invitrogen | C10640 |
| Formaldehyde, 16% (w/v), methanol-free | PierceTM | 28908 |
| Glass bottom dish, 35 mm | ibidi | 81218-200 |
| Poly-L-lysine, 0.01 % | Sigma | 25988-63-0 |
| PowerUp SYBR Master Mix | Thermo Fischer | A25742 |
| PrimeScript™ RT Master Mix | TaKaRa | RR036A |
| RNeasy Micro Kit | Qiagen | 74004 |
| Vectashield | Vector Labs | H-1000 |

### mRNA sequencing and data processing

mRNA sequencing (mRNA seq) libraries were generated from total RNA samples, isolated as stated above, by the Functional Genomics Centre Zurich (FGCZ) according to the TruSeq stranded mRNA library preparation protocol (Illumina). The NovaSeq 6000 instrument (Illumina) was used by the FGCZ to perform paired-end sequencing at 150 bp read length.

mRNA sequencing (mRNA seq) libraries were generated from total RNA samples, isolated as stated above, by the Functional Genomics Centre Zurich (FGCZ) according to the TruSeq Stranded mRNA library preparation protocol (Illumina). The NovaSeq 6000 instrument (Illumina) was used to perform paired-end sequencing at 150 bp read length for read 1 and read 2.

Raw .fastq files were processed using the RNA-seq analysis pipeline from snakePipes (61) with default parameters. Reads were aligned to the *Drosophila melanogaster* reference genome and the transcriptome reference annotation (Ensembl assembly dm6, release-94) using HISAT2 (62). Gene level quantification was performed using featureCounts (63) and for the genes in each pairwise comparison (Dorsal vs. lacZ; Ras^V12^, lacZ *vs*. lacZ and Ras^V12^, Dorsal *vs*. Ras^V12^, lacZ), differential expression analysis was performed using DESeq2 (64). Exploration of differentially expressed genes was done using the RNA-seq data visualization pipeline Searchlight 2 (v2.0.0) (65). The list of differentially expressed genes was adjusted using the cut-off p.adj < 0.05 and absolute log2fold > 0.5. The Gene Ontology (GO) and Kyoto Encyclopaedia of Genes and Genomes (KEGG) libraries were obtained from FlyEnrichr. RNA sequencing data generated in this study have been deposited in NCBI’s Gene Expression Omnibus (GEO) (66) and will be accessible through the GEO Series accession number GSE266879 (https://www.ncbi.nlm.nih.gov/geo/query/acc.cgi?acc=GSE266879).

## Acknowledgements

We would like to thank all current and former members of the Basler group, especially Vivien Fuchs, Jamie Little, Erich Brunner and Giulia Moro, as well as Hugo Stocker for fruitful discussions and advice. We also gratefully acknowledge Tor Erik Rusten and Caroline Dillard for their open communication during the completion of the manuscript.

This work, in particular FB was supported by the research grant Krebsliga Schweiz (no. KFS-4835-08-2019, https://www.krebsliga.ch/) and FB and FG are supported by the Swiss National Science Foundation (no. 207594, http://www.snf.ch/de/Seiten/default.aspx). FB was supported by the Candoc Forschungskredit (no. FK-20-085, https://www.research.uzh.ch/de/funding/phd/uzhcandoc.html).

---

**Fig S1.**
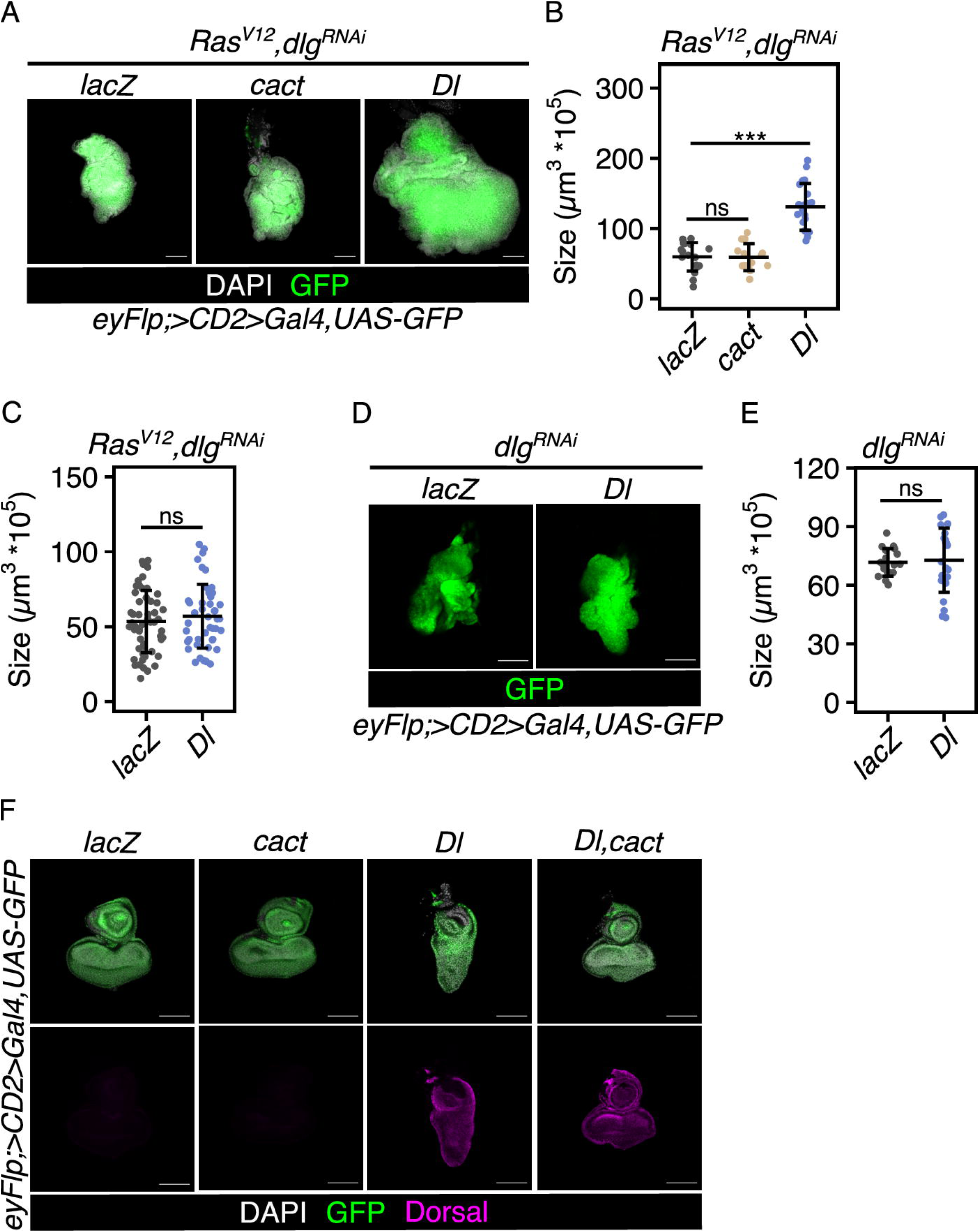
Toll signaling cooperates with oncogenic Ras to induce overgrowth. (A) Confocal images of Ras^V12^, dlg^RNAi^-induced EAD tumors of indicated genotypes at 120 h AED representative for tissue sizes quantified in (B). (B) and (C) Quantification of Ras^V12^, dlg^RNAi^-induced tumor volume of indicated genotypes at 120 h AED (B) and 96 h AED (C). Mean volume and standard deviation (error bar) are shown (***p < 0.001, ns: p = 0.99 (B) or ns: p = 0.41 (C), One-way ANOVA with post-hoc Tukey HSD). (D) Confocal images of dlg^RNAi^-transformed EADs of indicated genotypes at 120 h AED representative for tissue sizes quantified in (E). Images are extracted singles planes of samples mounted for 3D quantification. (E) Quantification of dlg^RNAi^*-*transformed EAD volume of indicated genotypes at 120 h AED. Mean volume and standard deviation (error bar) are shown (ns: p = 0.8, One-way ANOVA with post-hoc Tukey HSD). (F) Confocal images of EADs of indicated genotypes at 96 h AED labelled for Dorsal (magenta) to illustrate the efficiency of overexpressing Cactus to block the morphological defects caused by Toll signaling activation by Dorsal overexpression. In all confocal images DAPI (grey) is used to visualize nuclei and GFP (green) labels cells co-expressing indicated UAS-transgenes. Scale bars represent 100 µm. ns: not significant, EAD: eye-antennal disc, AED: after egg deposition, cact: Cactus, Dl: Dorsal, eyFlp: eyeless-driven Flippase.

**Fig S2.**
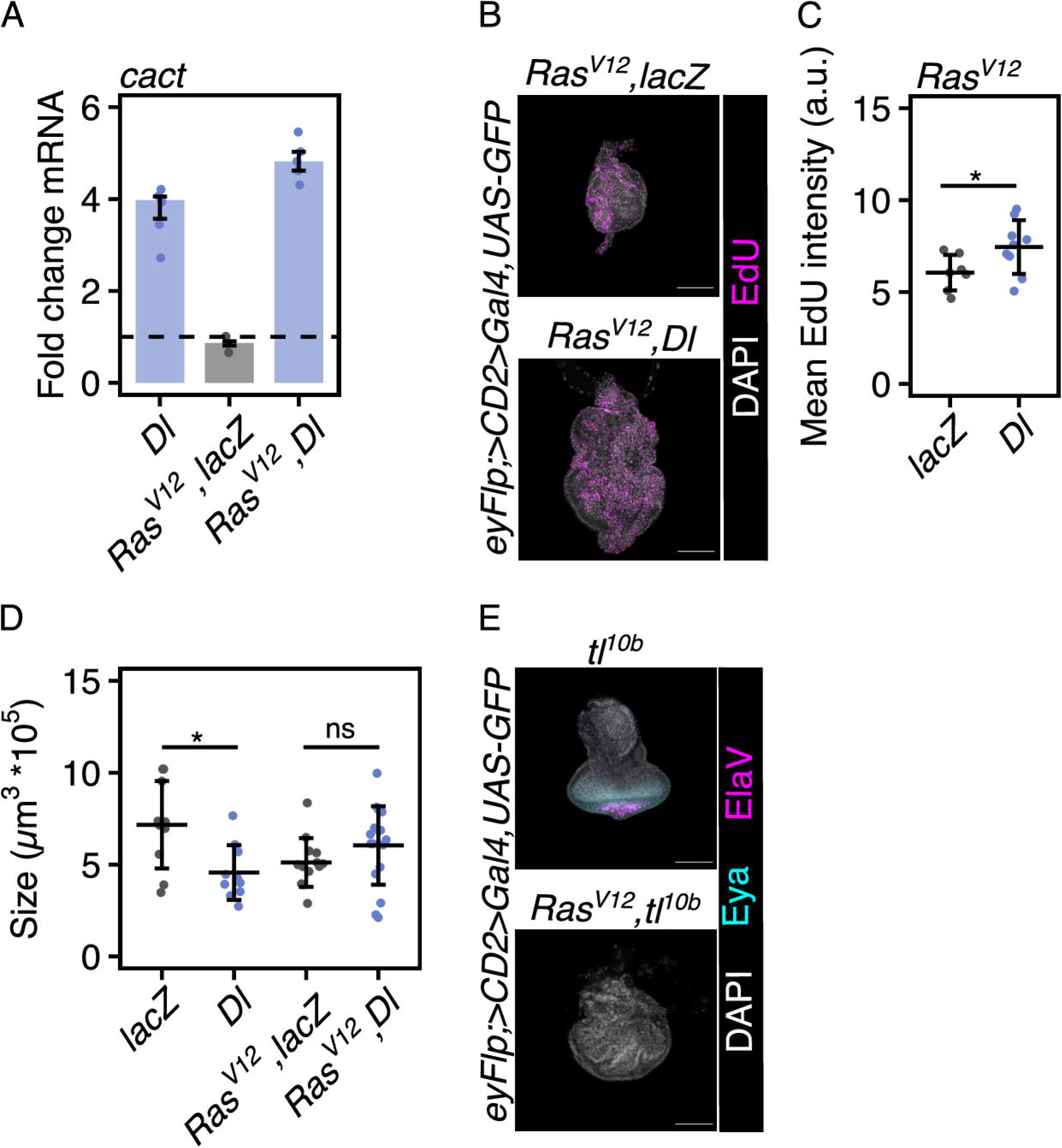
Dorsal inhibits retinal differentiation. (A) qRT-PCR analysis of *cact* mRNA levels in EADs expressing indicated UAS-transgenes under the control of an eye specific Gal4 (*eyFlp; act>CD2>Gal4*) as a read-out for Toll signaling activity. Data is shown as fold changes relative to the control (lacZ). n=3 biologically independent samples were analysed. Median fold changes and interquartile ranges (error bars) are shown. The dashed horizontal line illustrates the reference fold change of 1. (B) Confocal images of EAD tumors of indicated genotypes at 96 h AED stained with Click-iT™ EdU Alexa Fluor™ 647 (magenta) to identify DNA synthesis-based proliferation representative for EdU intensity quantified in (C). (C) Quantification of EdU mean fluorescence intensity per EAD tumor at 96 h AED. Mean intensity and standard deviation (error bar) are shown (*p < 0.05, One-way ANOVA with post-hoc Tukey HSD). (D) Quantification of tissue volume of indicated genotypes at 72 h AED illustrating the time-dependency of Dorsal-induced tumor overgrowth. Mean volume and standard deviation (error bar) are shown (*p < 0.05, ns: p = 0.55, One-way ANOVA with post-hoc Tukey HSD. (E) Confocal images of an EAD and tumor of the indicated genotype at 96 h AED labelled for retinal determination and photoreceptor differentiation markers Eya (cyan) and ElaV (magenta). In all confocal images DAPI (grey) is used to visualize nuclei and GFP (green) labels cells co-expressing indicated UAS-transgenes. Scale bars represent 100 µm. ns: not significant, a.u.: arbitrary unit. EAD: eye-antennal disc, AED: after egg deposition, Dl: Dorsal, tl^10b^: Toll^10b^, eyFlp: eyeless-driven Flippase, Eya: eyes absent, ElaV: embryonic lethal abnormal vision, EdU: 5-ethynyl-2’-deoxyuridine.

**Fig S3.**
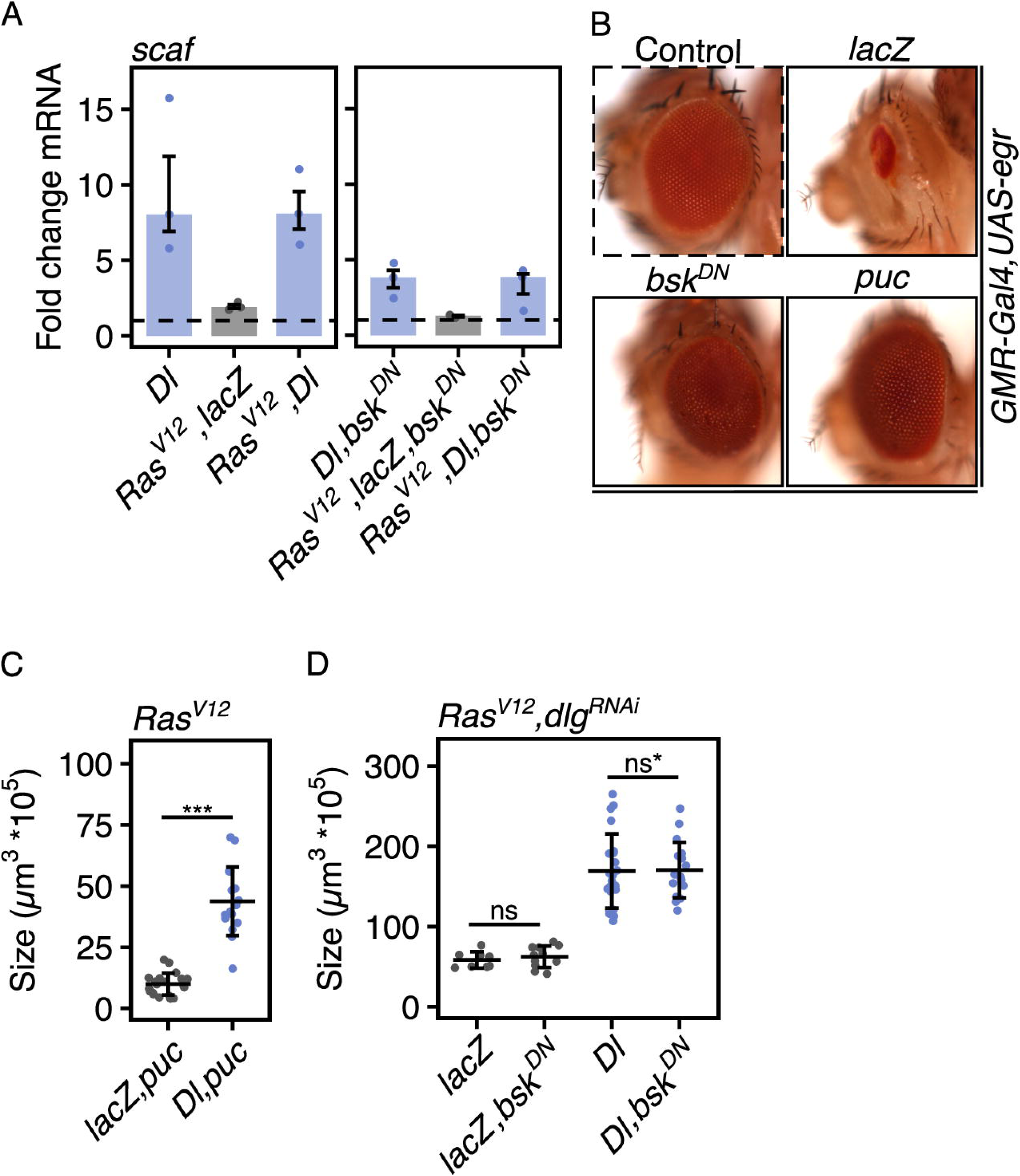
Overgrowth of Ras^V12^, Dorsal and Ras^V12^, dlg^RNAi^, Dorsal*-*induced tumors is independent of JNK signaling. (A) qRT-PCR analysis of *scaf* mRNA levels in EADs expressing indicated UAS-transgenes under the control of an eye specific Gal4 (*eyFlp; act>CD2>Gal4*) as exemplary confirmation of the JNK-dependent upregulation of target genes after overexpression of Dorsal. Data is shown as fold changes relative to the control (lacZ or lacZ,bsk^DN^). n=3 biologically independent samples were analysed. Median fold changes and interquartile ranges (error bars) are shown. The dashed horizontal line illustrates the reference fold change of 1. (B) Images of adult eyes of indicated genotypes showing the efficiency of overexpressing Bsk^DN^ or Puc to inhibit JNK signaling and rescue Egr induced loss of photoreceptor cells. All flies, except for wild-type control flies expressed GMR-Gal4, UAS-egr. (C) Quantification of tumor sizes of indicated genotypes at 96 h AED confirming the observation of JNK-independent induction of tumor overgrowth after Dorsal overexpression using co-expression of Puc to inhibit JNK signaling. Mean volume and standard deviation (error bar) are shown (***p < 0.001, One-way ANOVA with post-hoc Tukey HSD). (D) Quantification of tissue volume of indicated genotypes at 120 h AED illustrating that growth of Ras^V12^, dlg^RNAi^ tumors did not depend on JNK signaling at 120 h AED. Mean volume and standard deviation (error bar) are shown (ns: p=0.56, ns*: p=0.92, One-way ANOVA with post-hoc Tukey HSD). In all confocal images DAPI (grey) is used to visualize nuclei and GFP (green) labels cells co-expressing indicated UAS-transgenes. ns: not significant, EAD: eye-antennal disc, AED: after egg deposition, Dl: Dorsal, mmp1: matrix metalloproteinase 1, ets21c: Ets at 21c, bsk^DN^: dominant-negative Basket, Puc: Puckered, GMR: glass multiple reporter, egr: Eiger, JNK: Jun-N-terminal Kinase pathway, eyFlp: eyeless-driven Flippase.

**Fig S4.**
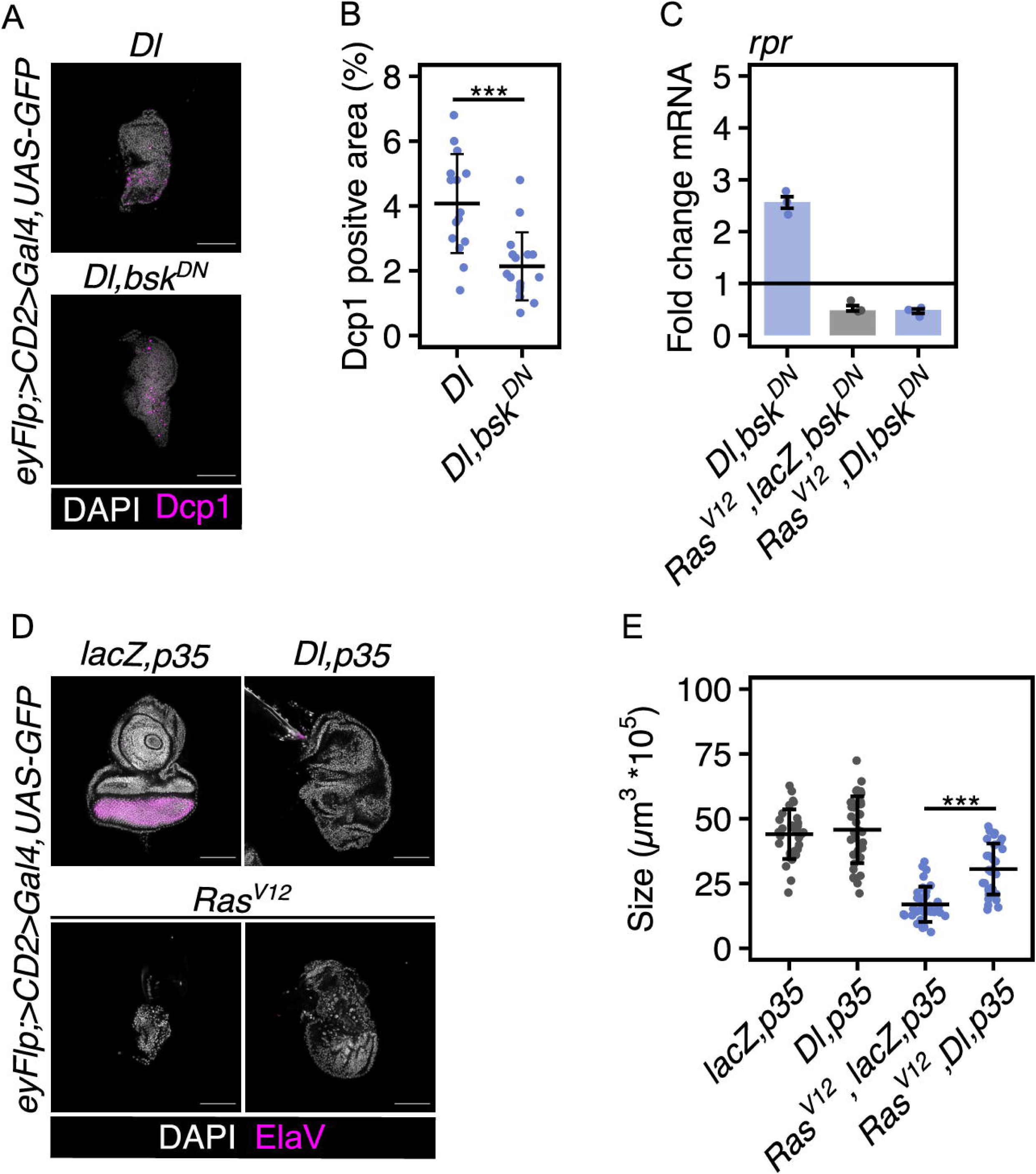
Dorsal-induced caspase-dependent cell death is suppressed by Ras^V12^. (A) Confocal images of EADs of indicated genotypes at 96 h AED labelled for Dcp1 showing the persistence of elevated levels of Dcp1 after JNK inhibition (B) Quantification of the Dcp1-positive area relative to whole disc size at 96 h AED comparing control EADs after overexpression of Dorsal with Bsk^DN^. Mean area and standard deviation (error bar) are shown (***p < 0.001, One-way ANOVA with post-hoc Tukey HSD). (C) qRT-PCR analysis of *rpr* mRNA levels in EADs expressing indicated UAS-transgenes under the control of an eye specific Gal4 (*eyFlp; act>CD2>Gal4*) illustrating the JNK-independent upregulation of pro-apoptotic gene expression after overexpression of Dorsal in control EADs. Data is shown as fold changes relative to the control (lacZ, bsk^DN^). n=3 biologically independent samples were analysed. Median fold changes and interquartile ranges (error bars) are shown. The dashed horizontal line illustrates the reference fold change of 1. (D) Confocal images of EADs of indicated genotypes at 96 h AED labelled for ElaV to highlight the lack of photoreceptor cells after Dorsal and p35 co-expression. (E) Quantification of tissue volume of indicated genotypes at 96 h AED to confirm the increased growth of discs overexpressing Dorsal after inhibition of apoptosis by co-expression of p35 using three-dimensional quantification. Mean volume and standard deviation (error bar) are shown (***p < 0.001, One-way ANOVA with post-hoc Tukey HSD). In all confocal images DAPI (grey) is used to visualize nuclei and GFP (green) labels cells co-expressing indicated UAS-transgenes. Scale bars represent 100 µm. ns: not significant, Dcp1: cleaved *Drosophila* caspase 1, JNK: Jun-N-terminal Kinase pathway, EAD: eye-antennal disc, AED: after egg deposition, Dl: Dorsal, Dcp1: cleaved *Drosophila* caspase 1, rpr: reaper, ElaV: embryonic lethal abnormal vision, bsk^DN^: dominant-negative Basket.

